# Efficient genome editing of human natural killer cells by CRISPR RNP

**DOI:** 10.1101/406934

**Authors:** Jai Rautela, Elliot Surgenor, Nicholas D. Huntington

**Author notes:** Corresponding Authors: Prof. Nicholas Huntington,; Dr. Jai Rautela.

## Abstract

The ability to genetically modify CD8 T cells using viral gene delivery has facilitated the development of next generation of cancer immunotherapies such as chimeric antigen receptor (CAR) T cells engineered to specifically kill tumor cells. Development of immunotherapies targeting natural killer (NK) cells have stalled in part by their resistance to viral gene delivery. Here, we describe an efficient approach to genetically edit human NK cells by electroporation and CRISPR-Cas9 ribonucleoprotein (RNP) complexes. We detail electroporation pulse codes and buffer optimization for protein uptake by human NK cells and viability, and the efficiency of this approach over other methods. To highlight the transformative step this technique will have for NK cell immunotherapy drug discovery, we deleted NKp46 and CIS in primary human NK cells and validated murine findings on their key roles in regulating NK cell anti-tumor function.

## Introduction

Immunotherapies are emerging treatment options for several cancer types with CD8 T cells being the primary therapeutic target. The efficiency of viral gene delivery to activated CD8 T cells *in vitro* has facilitated the clinical development of chimeric antigen receptor (CAR) T cell recognizing tumor antigens such as CD19 on B cell leukemia and have recently been approved by regulatory agencies. Natural killer (NK) cells are an additional anti-tumor effector population with a clear role in limiting tumor metastasis^1,2^ and an emerging role in orchestrating inflammation and immune infiltration within solid tumors^3,4^. NK cells are dependent on the cytokine IL-15 for their development and maturation^5^, and are educated in the periphery through the signals transduced through an array of germline activating and inhibitory receptors^6^. In addition to numerous soluble factors, it is the balance of signals transduced through these receptors that also dictates whether an NK cell engages in a cytotoxic attack against a target cell. We previously described CIS (encoded by *Cish*) as a potent checkpoint in NK cell activation by limiting IL-15 signal transduction^7^, while CBL-B (*Cblb*) has also been proposed as an intracellular repressor of NK cell activation^8^. In addition, the group of Mantovani described IL-1R8 (*Il1r8*) acting as a negative regulator of IL-18 responsiveness in NK cells^9^ while more recently, TIGIT (*Tigit*) was shown to be a major inhibitory receptor on NK cells^10^ with deletion of *Cish, Cblb, Il1r8* or *Tigit* resulting in dramatically improved NK cell-dependent anti-tumor immunity.

These pre-clinical findings highlight that NK cell anti-tumor immunity can be augmented *in vivo* and as such, development of NK cell immunotherapy drugs that target such pathways should be investigated. In contrast to CD8 T cells, NK cells are yet to be specifically or efficiently targeted by immunotherapy drugs in human cancer^6,11^. A major hurdle to NK cell drug discovery has been target validation in human NK cells and this is indeed an important step given the clear disparities in NK cell phenotypes and frequencies between species. The field has therefore awaited the development of rapid and efficient gene-editing technologies such as CRISPR-Cas9 in order to perform targeted gene-editing to understand the biology on primary human NK cells. Since the initial description of CRISPR-Cas9 in mammalian cells, this technology has been used to great effect to edit genes in several immune cell lines and more recently, primary human T cells^12,13,14^. Editing immune cell lines has been relatively simple through lentiviral delivery of Cas9 and a gene-specific guide RNA, and has allowed genome-wide screening studies to be performed^15^. The efficacy of using similar viral delivery systems in primary immune cells has been highly cell-type specific. NK cells are notoriously difficult to infect with retro/lentiviral particles meaning that tried and tested genetic modification approaches for human T cells are not appropriate for human NK cells. For example, while IL-2 or IL-15 results in robust activation and proliferation of human NK cells, typically only around % of these cells will be infected with a lentivirus. When lentiviral multiplicity of infection (MOI) was increased from 1 to 20, the efficiency of lentivirus transduction on human NK92 cells was unchanged^16^, while other groups have increased the MOI to 150 and still only achieve ~10% transduction of human NK cells^17^. Electroporation of NK cells with mRNA or plasmid DNA is efficient for transient expression (such as anti-CD19 CAR) yet there is no evidence that transient Cas9 and sgRNA expression can lead to efficient gene editing in NK cells^17,18^. While viral delivery of the Cas9 system to primary cells is the most utilized approach at present, delivery of recombinant Cas9 in complex with a gene-specific gRNA (together termed a ribonucleoprotein, RNP) is an emerging alternative. Reports have demonstrated that delivery of this complex can result in gene-editing *in vivo*^19^ and highly efficient in primary immune cells, including human T cells^12,13,14^. With our extensive experience with primary human NK cells, we set out to firstly establish a robust and efficient DNA and viral-free CRISPR-RNP gene-editing strategy for primary human NK cells, and secondly, use this technical breakthrough to validate our recent discovery of an NK cell checkpoint in primary human NK cells. The efficiency and simplicity of our CRISPR-RNP gene-editing strategy will expedite target validation in NK cells and be transformative for immunotherapy drug development.

## Methods

### Cell culture & lentivirus production

Primary human NK cells were isolated from peripheral blood samples using an immunomagnetic negative selection kit (Stemcell Technologies, #17955) and expanded in G-Rex plates (Wilson Wolf, 80240M) using NK-MACS media (Miltenyi, 130-114-429) supplemented with 5% AB human serum (Sigma-Aldrich, H6914), 1000 IU.mL of human IL-2 and 20 ng.mL human IL-15 unless otherwise stated. The SNK6 and SNK10 cell lines were a gift from (Professor Maher Ghandi, The University of Queensland) were grown in NK MACS media supplemented with 10 ng.mL human IL-15. The Daudi Burkitt’s lymphoma cell line were maintained in RPMI containing 10% FBS. Cas9-containing lentivirus was produced according standard procedures using the FUCas9-mCherry plasmid described in^20^. Fresh lentivirus was used to spin-infect human T and NK cell lines on two consecutive days.

### CRISPR protocols, reagents & sequencing

#### Optimization of electroporation conditions

Expanded primary human NK cells (500,000 cells per well of a 16-well Nucleocuvette strip) were used to sequentially optimize the pulse code and buffer conditions for maximum viability and uptake of 70kDa FITC-labelled dextran (Sigma #46945). A range of custom buffers were created and tested, with Solution 2 + mannitol (5mM KCl, 15mM MgCl_2_, 15mM HEPES, 150mM Na_2_HPO_4_/NaH_2_PO_4_ (phosphate buffer), 50mM mannitol, pH 7.2) resulting in the best combination of viability and dextran-uptake (see RNP protocol below for electroporation handling technique). All electroporation experiments were carried out using the 4D Nucleofector system (Lonza).

### Evaluation of the transient Cas9 expression system

Primary human NK cells (isolated and rested overnight) and human NK cell lines were electroporated using the optimized electroporation conditions (CM137 & solution 2+mannitol) with a positive control plasmid (pmaxGFP, Lonza), dextran-FITC or a px458 vector encoding spCas9 and a sgRNA against human CD45 (vector and cloning described by^21^, sgRNA sequence in table below). Transfection efficiency (dextran), transcription/translation efficiency (pmaxGFP) and CD45-deletion efficiency (px458-CD45) were assessed by FACS 60 and 144 hours after electroporation (see RNP protocol below for electroporation handling technique).

#### Evaluation of Cas9-RNP system

Primary human NK cells (isolated and rested overnight) and human NK cell lines were electroporated using the optimized electroporation conditions (CM137 & solution 2+mannitol) with Cas9-RNP complexes targeting human CD45. Deletion efficiency was assessed 120 hours post-electroporation, and various SNK10-CD45 populations were FACS sorted and cultured for 14 days to show stability of the phenotype. The optimized RNP workflow follows:

1. Prepare cells for Electroporation
  a. Pre-warm culture media to 37°C
  b. Pellet 500,000 cells primary NK/NK cell lines by centrifuging at 400 x g for 5 minutes at room temp
2. Form the Cas9/gRNA complex
  a. Cas9 nuclease provided at concentration of 61μM (IDT, 1081059)
  b. Resuspend electroporation enhancer to 100μM in TE buffer (IDT, 1075916)
  c. Resuspend sgRNA in to 100μM in TE buffer (Synthego)
  d. Mix components as follows (for one well of a 16-well Nucleocuvette strip)
    i. 5μl 100μM sgRNA (or molar equivalent if using crRNA:tracrRNA 2-part system)
    ii. 1.7μl 61μM Cas9
    iii. 1μl 100μM enhancer
  e. Incubate at room temperature for 10 minutes
  f. Electroporation
    a. Aspirate supernatant from centrifuged cells
    b. Gently resuspend cells in 13μl electroporation buffer (Sol2 + mannitol for human NK) and mix with 6.7μl of the RNP mixture (total of approx. 20μl), avoid getting bubbles
    c. Transfer entire mixture to one well of 16-well Nucleocuvette strip
    d. Gently tap the 16-well strip to ensure no air bubbles (or use needle/pipette to remove any big bubbles)
    e. Select wells for electroporation and specify pulse code (CM137) and buffer used (set buffer ‘P3’ as a surrogate for solution 2 + mannitol)
  g. Post-electroporation handling
    a. Very gently add 100μl of pre-warmed complete media to each well of the 16-well strip and incubate at 37°C/5% CO_2_ for 10 minutes
    b. Very gently transfer cells to appropriate flask/plate, avoiding excessive pipetting

#### Application of Cas9-RNP system

Primary human NK cells (isolated and rested overnight) and human NK cell lines were electroporated using the optimized electroporation conditions above with Cas9-RNP complexes targeting *NCR1*, or *CISH*. Cells were allowed to recover and expand for 5 days after electroporation in media containing 5ng.ml IL-15 and deletion efficiency was analyzed by western blotting, FACS or next-generation sequencing. For studies targeting *NCR1*, the NCR1^+^ and NCR1^*-*^ populations were subsequently FACS sorted and used in cytotoxicity assays. Next generation sequencing of CIS indel frequencies was performed using standard NGS protocols, briefly; PCR reactions containing sample gDNA and primers (listed below) flanking the CRISPR binding site were used to produce amplicons that were subjected to a second round of PCR to introduce indexes and sequencing adaptors. These ‘indel libraries’ were then purified (AMPure XP, Beckman Coulter), pooled for multiplexing, and sequenced on an miSeq instrument (illumina) with a target of 10-20,000 reads.

sgRNA and primer sequences used in this study:

Target: Guide# Guide sequence (5’->3’)

*PTPRC* 61 AGTGCTGGTGTTGGGCGCAC

*NCR1* 543 TGGGGCTCGGCCCAGATGAA

*NCR1* 544 TCTCCCAAAACCGTTCATCT

*CISH* 26 CTCACCAGATTCCCGAAGGT

*CISH* 57 CCGCCTTGTCATCAACCGTC

*CISH* NGS primers: Fwd: AGAGAGTGAGCCAAAGGTGC, Rev: GGGCTATGCCTCCCAAATCA

### Protein assays & Flow cytometery

Flow cytometry antibodies and reagents: CD335 (NKp46; clone REA808 Miltenyi Biotec), CD45 (clone REA747 Miltenyi Biotec), human NKp30/NKp46-Fc reagents were a kind gift from Prof. Ofer Mandelboim (Hebrew University, Israel).

Western blotting reagents: JAK1 (B-3, #sc-376996 Mouse monoclonal, Santa Cruz, 130kDa), Phospho-JAK1 (Tyr1022, Tyr1023, #44-422G, Rabbit polyclonal, Thermo Fisher, 130kDa), Phospho-STAT5A/B (Tyr694/699, 8-5-2, #05-495, Mouse monoclonal, Merck, 90kDa), STAT5A (ST5a-2H2, #13-3600, Mouse monoclonal, Thermo Fisher, 90kDa), CISH (D4D9, #8731S, Rabbit monoclonal, Cell Signaling Technology, 32kDa, 38kDa), Actin (I-19, #sc-1616, Goat polyclonal, Santa Cruz 45kDa). Samples were processed using RIPA lysis buffer (50mM Tris pH 7.4, 150mM NaCl, 0.25% deoxycholic acid, 1% NP-40, 1mM EDTA, Millipore) containing cOmplete protease inhibitors (Roche), PMSF (#8553, Cell Signaling Technology), sodium orthovanadate (NEB) and PhosSTOP (Roche). Protocol: cell pellets (1.2 million) were lysed in 60 μl lysis buffer with protease inhibitors for 30m ice and centrifuged at 13,000 x g for 5 minutes to clear cell debris. 10 μl of 6X sample buffer was added to 50 μl of clarified lysate, incubated at 85°C for 5 minutes and 30 μl was run on a 4-12% gel using MOPS buffer. Primary antibodies were used at a 1/1000 dilution in PBS containing 5% BSA, 0.1% Tween-20 and incubated overnight at 4°C. Secondary antibodies were used at 1/3000 in the same diluent and incubated for 1 hour at room temperature.

### Cytotoxicity & proliferation assays

NK cell cytotoxicity assays were performed as previously described^22^. Briefly, target cells were loaded with calcein-AM, mixed with NK cells at various ratios, incubated for 4 hours at 37°C and analyzed by transferring 100 μl of cell-free supernatant into opaque 96-well plates and comparing the fluorescent intensity of test wells to positive (target cells + 2% triton) and negative (target cells alone) controls. For proliferation assays, cells were plated at the indicated starting densities and absolute live cell number was quantified daily using FACS by adding enumeration beads (123count eBeads, Thermo Fisher) and a viability dye (propidium iodide).

### Humanized mice

Human cord blood cells obtained from the Bone Marrow Donor Institute/Cord Blood Research Unit (Murdoch Children’s Research Institute, Australia) following approval from The Walter and Eliza Hall Institute of Medical Research Human Research Ethics Committee. CD34^+^ HSCs were isolated from fresh human cord blood cells using a CD34 positive-selection kit (Stemcell technologies, #17896) and cultured in using in StemMACS HSC expansion media (130-100-473, Miltenyi) supplemented with StemMACS HSC expansion cocktail (130-100-843, Miltenyi) for 3 days. On the third day, CD34 cells were cleaned-up using Ficoll-Paque PLUS (GE Life Sciences) density centrifugation to remove dead cells and magnetic particles, and then electroporated (using the same conditions optimised for primary NK cells) with *CISH* g26-RNP or mock RNP. Cells were returned to culture for a further 2 days to recover, before being injected into the facial vein of 2 day old sub-lethally irradiated BRGS (BALB/c Rag2^tm1Fwa^ IL-2Rγ^tm1Cgn^ SIRPα^NOD^ ^Flk2tm1Irl^) mice^23^. Mice were allowed to reconstitute for 12-15 weeks before being analysed for human NK cell development. Splenic NK cells were also FACS sorted and used in a proliferation assay (seeded at 10,000 cells/well in 96-well round-bottom plates) with various concentrations of human IL-15. On the final day of the proliferation assay, the well containing *CISH* g26-RNP cells in 5ng.mL IL-15 were harvested and subjected to NGS to determine the relative number of indels in the CISH gene.

## Results

### Optimization of electroporation and protein delivery into primary human NK cells

The growing applications for adoptive T cell therapies stems from the relative ease of introducing and stably expressing synthetic genes via retro/lentiviral transduction. Genetic modification of T cells via CRISPR also exploits this ease of transduction. To compare the efficiency of human NK cell viral transduction versus human T cell transduction and the potential for genetic modification via CRISPR/Cas9, we transduced human NK (NKL) or T (Jurkat) cell lines with FUCas9-mCherry lentiviral particles (MOI = 5). Both cell lines were actively dividing *in vitro*, and two rounds of standard spin-infection were used. Consistent with previous work, only T cells, not NK cells were efficiently infected and stably expressed the viral gene products Cas9-mCherry (**Fig. 1a**). To investigate alternate approaches to genetically modify human NK cells we next developed a feeder-free method for primary human NK cell expansion in order to optimize electroporation conditions on a consistent source of material. NK cells are frequent in the blood and their isolation can be achieved by numerous methods including positive selection by magnetic beads and FACS sorting. We opted for a negative selection approach as this typically yields >95% purity (NKp46^+^CD56^+^CD3*-*) with minimal loss of viability as is often associated with FACS sorting. NK cells were then seeded at a density of 2e6 cells.cm^2^ in either G-rex 24 or 6 well plates using NK-MACS media (Miltenyi) supplemented with 1000 IU.mL IL-2, 20ng.mL IL-15 and 5% AB+ human serum. Half the culture media was removed and replaced with fresh media every third day until desired number of NK cells was achieved (**Fig. 1b**). Numerous methods exist to deliver small molecules and nucleic acids into cells, however delivery of large protein complexes is most efficient when the cell surface is physically disrupted to allow the passive diffusion of such molecules into the cells. Both mechanical disruption and electroporation-based technologies have been adapted to introduce CRISPR RNPs into mammalian cells, though electroporation has emerged as the predominant platform due to the widespread availability of these devices and adaptability for use in in-vivo settings^19^. However, one of the major hurdles in the NK cell field has been the relatively poor survival of NK cells post-electroporation. We therefore next establish electroporation conditions for human NK cells that would maximize their viability and uptake of large molecules using the widely-available Lonza 4D nucleofector device (**Fig. 1b**). Beginning with the manufacture’s recommendations, we set out to optimize the electrical pulse applied to the cells, or “pulse-code”, using the manufacturer’s own P3 nucleofection buffer (**Fig. 1c**). The recommended read-out for such optimizations is the nucleofection of a GFP-encoding plasmid to determine the efficacy of nucleofection. However, given the size of recombinant Cas9 (approximately 160kDa) and the fact that RNP-mediated editing does not require transcription from a plasmid template we decided to optimize with a fluorescently-labelled high molecular weight dextran. Pulse code DN100 resulted in the maximum up-take of FITC-dextran (~95%), yet viability was only ~40% (**Fig. 1c**). We next tested whether alternate buffers could further improve NK cell viability and therefore created a panel of home-made buffers to compare to the P3 standard. Our home-made buffer “Sol. 2 + mannitol” improved NK cell viability (>60%) without compromising FITC-Dextran uptake and was therefore adopted as our standard buffer to for primary NK cell electroporation (**Fig. 1d**). We then re-optimized the pulse-codes using our Sol. 2 + mannitol buffer and found that pulse-code CM137 was optimal resulting in >80% NK cell viability and FITC-Dextran uptake and Sol. 2 + mannitol/pulse-code CM137 was adopted as our human NK cell electroporation conditions (**Fig. 1d**).

**Figure 1.**
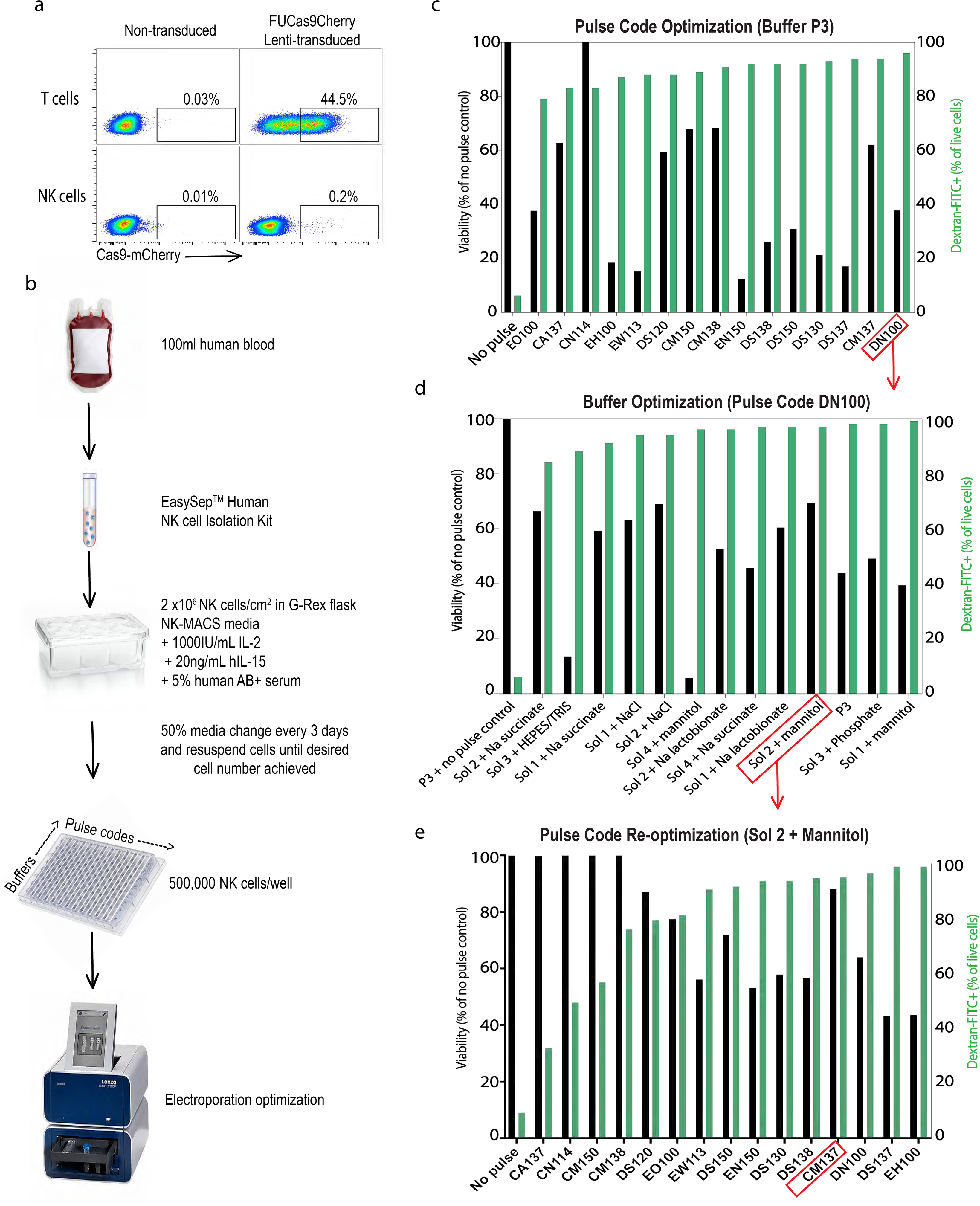
The unmet need of efficient gene editing of human NK cells. (**a**) Human T cell (Jurkat) and NK cell (NKL) lines were spin-infected with FUCas9Cherry lentiviral particles (MOI = 5) and Cas9-mCherry monitored by FACS at 72 hours. Data are representative of 3 independent experiments. (**b**) Flow chart of human NK cell isolation, expansion and electroporation optimization. Lonza 4D electroporation optimization for primary human NK cells; (**c**) Pulse code optimization for using buffer P3, (**d**) Buffer optimization using DN100 pulse code and (**e**) Pulse code re-optimization using Sol. 2 + Mannitol buffer. Black bars indicate NK cell viability, green bars indicate uptake of FITC-dextran (electroporation efficiency). Data are representative of 2 independent experiments.

### Transient transfection of Cas9 RNP but not Cas9-expressing plasmid results in efficient gene editing in human NK cells

A recent report demonstrated that T cells could be efficiently gene-edited by simply introducing plasmids that contain Cas9 and the gRNA (transient transfection)^24^, and if applicable to NK cells, would represent the most rapid and cost-effective method of gene-editing. We therefore cloned one of our validated guides against human CD45 (guide #61) into the Px458 ‘all-in-one’ vector^21^ that encodes Cas9 gene, eGFP and the sgRNA. A vector encoding eGFP alone (pmaxGFP) was used as a positive control for transient gene expression. 60 and 144 hours following electroporation of primary human NK cells with these plasmids, we measured NK cell expression of GFP and CD45 (targeted for deletion by PX458). While primary human NK cells were found to expressed GFP (25-35%) following electroporation of pmaxGFP, we failed to detect GFP expression using Px458 and therefore unsurprisingly no deletion of CD45 was observed (**Fig. 2a**). The fraction of GFP expressing cells was significantly higher (50-80%) when human NK cell lines (SNK6, SNK10) where electroporated with pmaxGFP, yet still no Cas9-GFP nor CD45 deletion was observed in these lines (**Fig. 2a**). Thus, we concluded that primary human NK cells poorly transcribe genes from plasmid templates, which is further diminished when required to transcribe large genes such as Cas9 in the case of Px458.

**Figure 2.**
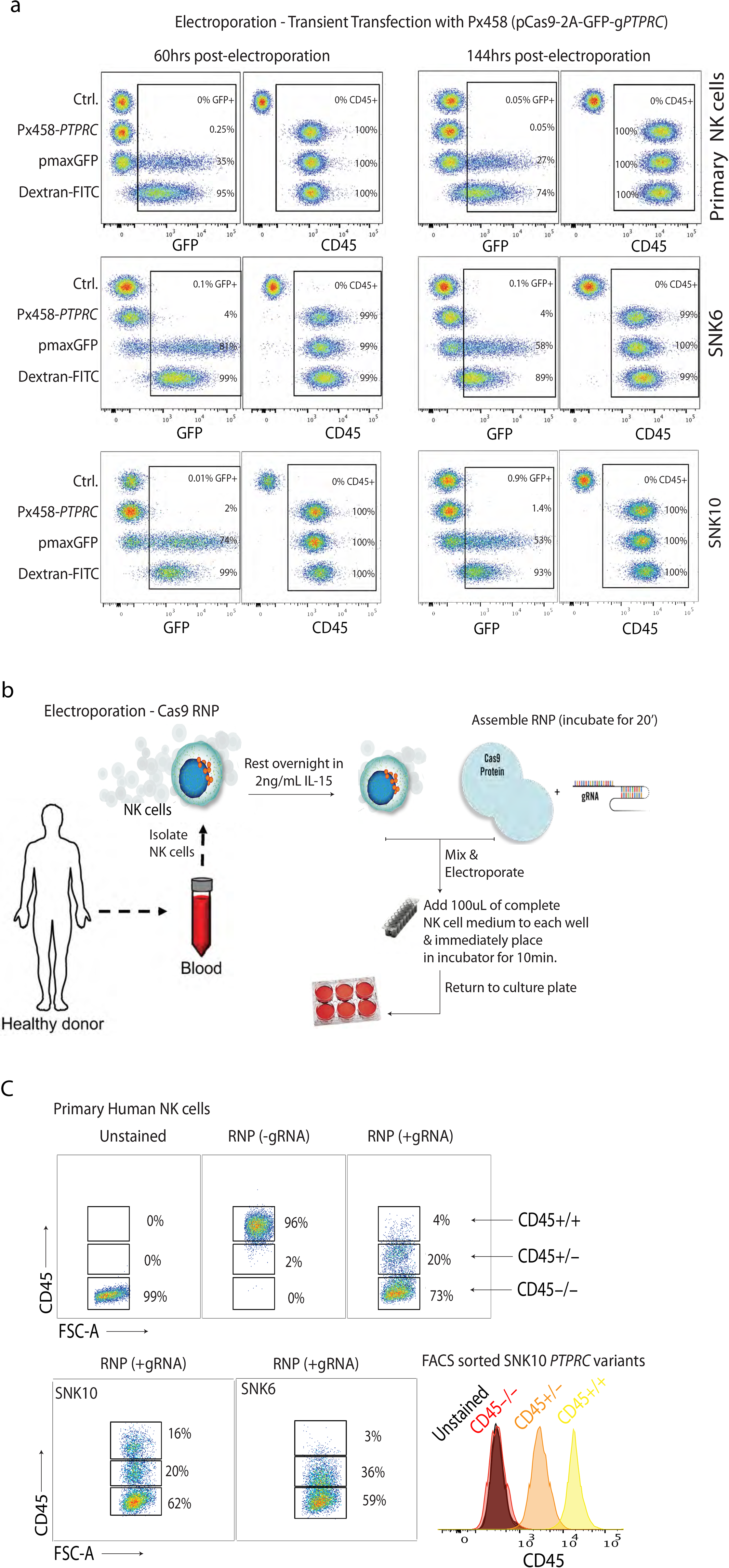
Cas9-RNP results in rapid, efficient gene deletion in primary human NK cells. (**a**) A transient Cas9 and sgRNA expression system (Px458) was tested for the ability to delete *PTPRC* (CD45) from NK cell lines (SNK6/SNK10) and primary human NK cells (dextran-FITC and pmaxGFP served as transfection and transcription controls, respectively). All data are shown. (**b**) Schematic of workflow for isolating and electroporating primary human NK cells. (**c**) Electroporation of Cas9 RNP complexes (carrying the same sg*PTPRC* sequence used in the px458 vector in (**a**) results in highly efficient deletion of CD45 from SNK10 and SNK6 human NK cell lines and primary human cells after 3-5 days in culture. Histograms show the stability of FACS-sorted CD45-expression variants (high, medium and low) in the SNK10 line after 2 weeks in culture. Data are representative of 2 independent experiments for SNK10/SNK6 and 4 individual healthy adult donors for primary NK cells.

Considering the difficulty of inducing Cas9 expression within NK cells, we opted for the delivery of the fully assembled “ready-to-go” ribonucleoprotein (RNP) complexes as this has recently been shown to efficiently edit primary human T cells (**Fig. 2b**)^13,14^. RNP complexes were created by incubating fully-synthetic sgRNA guides (identical in sequence to those used in the Px458 plasmid) with recombinant Cas9 prior to electroporation into *in vitro* expanded primary human NK cells as in **Fig. 1**. NK cells were then rested in culture for 3-5 days to allow sufficient time for protein turnover. By flow cytometry, the extent of gene editing after this single-step, single-guide, manipulation appeared to be remarkably efficient. In both primary human NK cells and NK cell lines (SNK10 and SNK6) we observed loss of both copies of *PTPRC* (CD45^-/-^) in 55-75% of the cells, whereas between 3-20% of the NK cells remained unedited (CD45^+/+^**Fig. 2c**). We also detected a population of cells (20-40%) which expressed intermediate levels of CD45 and likely represents heterozygous deletion *PTPRC* (CD45^+/-^). We next FACS sorted SNK10 cells based on CD45 expression (high, intermediate, negative) 5 days after Cas9-RNP was performed. SNK10 CD45 variants were maintained in culture for 2 weeks and CD45 expression was found to be stable (**Fig. 2c**) suggesting homo- and heterozygous human NK cells can be generated and that Cas9-RNP is an efficient method to genetically modify large numbers of primary human NK cells.

### Using Cas9 RNP to validate genetically-modified murine NK cell data in human NK cells

The NCR-family of NK cell activating receptors includes NKp46, NKp44, NKp30 (encoded by *Ncr1, Ncr2, Ncr3*) in humans but only NKp46 exists in mice^25^. NKp46 is ubiquitously expressed on mature NK cells where it defines the lineage in both species and can trigger IFNγ production when cross-linked^7^. NKp46-null murine NK cells (*Ncr1*^*gfp/gfp*^) display impaired ability to kill NKp46-ligand expressing tumor cells *in vitro* ^26,27^. To investigate if optimal human NK cell cytotoxicity against tumor cells also requires NKp46, we deleted *NCR1* from freshly isolated primary human NK cells from two independent donors using Cas9-RNP with two distinct gRNA against *NCR1*. Deletion efficiency ranged from 8-23%, with both gRNA having similar *NCR1* deletion efficiency (loss of NKp46 expression; **Fig. 3a**). We next FACS sorted NKp46^+^ and NKp46-NK cells from each donor and directly compared their cytotoxicity against Daudi cells (human Burkitt B cell lymphoma). Daudi cells were chosen due to their susceptibility to lysis by allo-NK cells and because they stained highly for human NKp46-Fc indicating they express putative NKp46 ligands (**Fig. 3b**). Consistent with the murine data, we found human NKp46^-^ NK cells were clearly inferior to NKp46^+^ counterparts from the same donor at lysing Daudi cells *in vitro* (**Fig. 3c**). Given the low deletion efficiency of NCR1 (% NKp46*-*) in freshly isolated primary human NK cells, we next investigated if highly metabolic, activated and proliferating primary human NK cells were a more efficient target population for *NCR1* deletion. NK cells isolated from donor 2 (**Fig. 3a)** were expanded in G-Rex flasks for 14 days as in **Fig. 1**. Using the identical g*NCR1*/g543 as in **Fig. 3a**, we observed significantly higher *NCR1*-deletion using expanded primary human NK cells with 85% NKp46*-* NK cells (5μl gRNA; **Fig. 3d**) compared to 23.7% NKp46*-* NK cells using freshly NK cells from the same donor (**Fig. 3a**). We also observed that *NCR1* gene-editing efficiency could be modulated by titrating the amount of gRNA used in the RNP complex and including a non-homologous DNA enhancer as previously shown^28^ (**Fig. 3d**). Taken together, these data support a conserved role for NKp46 in augmenting NK cell cytotoxicity upon detection of NKp46-ligands on tumor cells and highlight the utility of Cas9-RNP in validating murine NK cell biology in humans.

### Validating *CISH* as an immune checkpoint in human NK cells using Cas9-RNP

Work from our group and others over the past 10 years has established the cytokine IL-15 is essential for most facets of NK cell biology including survival, homeostasis, differentiation and activation^1,7,29^. We recently discovered that CIS (encoded by *Cish*) acts as a checkpoint in mouse NK cell activation by negatively regulating IL-15 signaling^7^. The heightened IL-15 responsiveness of *Cish*^*-/-*^ NK cells rendered *Cish*^*-/-*^ mice resistant to melanoma, prostate and TNBC metastasis. A hallmark of *Cish* loss-of-function in murine NK cells is hyper-responsiveness to IL-15 *in vitro* which is read-out as enhanced proliferation, enhanced phospho-JAK1 and phospho-STAT5 thus we next generated *Cish*-null human NK cells to investigate these parameters. Firstly, we targeted *CISH* for deletion in SNK10 and SNK6 human NK cell lines to test the efficiency of the gRNA. Using two independent *Cish* gRNA (g26 and g57) or pooled *CISH* gRNA (g26 + g57) resulted in a loss of CIS protein in both SNK6 and SNK10 with effiency being 80-90% based of frequency of insertion/deletion mutations (indels) in *CISH* determined by Next-Gen Sequencing (NGS; **Fig. 4a and data not shown**). Both single g26 and g57 showed similar efficiency for *CISH* deletion and SNK10 cells electroporated in the absence of Cas9-RNP (SNK10-Mock) served as negative controls. We next probed SNK10+g57 cells for IL-15 hyper-responsiveness compared to SNK10-Mock. Both phospho-JAK1 and phospho-STAT5 were readily induced by IL-15 from 15 minutes, indicating the IL-15 pathway is intact in SNK10 cells (**Fig. 4b**). Consistent with our murine findings in *Cish*^*-/-*^ NK cells, SNK10+g57 cells displayed clearly enhanced phospho-JAK1 and phospho-STAT5 following IL-15 stimulation compared to SNK10-Mock. CIS protein was induced in SNK10-Mock cells at 60 minutes following IL-15 stimulation, consistent with murine NK cells and the ~30kDa CIS protein was absent from SNK10-g57 cells while a weak ~35kDa CIS protein band could be detect at 120 minutes inline with the ~10% unedited cells in the pool (**Fig. 4b**). As previously documented for *Cish*^*-/-*^ murine NK cells, *in vitro* culture in 5ng.mL of IL-15 resulted in significantly greater SNK10-g57 cell proliferation compared to SNK10-Mock controls (**Fig. 4c**). We next confirmed these findings in primary human NK cells using the identical protocol. NK cells from peripheral blood of 4 healthy human donors were isolated, cultured in IL-15 overnight and electroporated in the presence of Cas9+g26 (g26-RNP) or g26 alone (Mock-RNP). NK cells were then cultured in 5ng.mL IL-15 for 5 days. In each of the 4 donors, g26-RNP resulted in a 2-3 fold increase in NK cell numbers compared to Mock-RNP from the same donor (**Fig. 5a**) and this was in part due to enhanced proliferation (**Fig. 5b**). Being an intracellular protein that is rapidly induced then degraded following IL-15 stimulation, CIS levels are difficult to quantitate using conventional techniques such as western blotting and intracellular FACS, and this is complicated further by the fact that two CIS protein isoforms (~30kDa/35kDa) are detected by the available anti-CIS polyclonal antibodies. g26-RNP results in loss of both CIS isoforms (**Fig. 4a**) therefore we used NGS to quantify the frequency of DNA mutation (indels) caused by Cas9+g26 around the *CISH* target sequence. A range of idels were introduced by g26-RNP in primary human NK cells with the average indel frequency being 63.75% and ranging from 45-80% indel frequency across the 4 donors (**Fig. 5c**). Furthermore, primary human NK cells edited by g26-RNP, g57-RNP or both all displayed enhanced cytotoxicity towards Daudi B lymphoma cells compared to Mock-RNP control cells (**Fig. 5d**) mimicking the enhanced murine *Cish*^*-/-*^ NK cell *in vitro* cytotoxicity towards allogeneic targets^7^. Despite their hyper-responsiveness to IL-15, *Cish*^*-/-*^ NK cells do not accumulate in *Cish*^*-/-*^ mice^7^. Our extensive experience with human xenograft models or humanized mice promoted us to investigate whether our Cas9-RNP protocol could be adapted to CD34^+^ cord blood progenitors to generate humanized mice lacking *CISH* in all human immune cells and validate *in vivo* safety of *CISH* deletion/inhibition on human NK lymphopoiesis. Again, we opted to perform these experiments using the single g26 so that indel frequency could be easily estimated with NGS throughout the experiment. Using the same electroporation conditions as primary human NK cells, we were able to achieve 43% editing efficiency in freshly-isolated human cord blood CD34^+^ cells. This result was encouraging for only a single guide and the fact that CIS is not highly expressed in CD34 cells, an obvious impediment to efficient gene editing in cells^30^. These pool of partially *CISH*-deleted/g26-RNP cells or Mock-RNP cells were then injected into the facial vein of lethally-irradiated 3-day old BRGS mice^23^ and allowed to reconstitute for 12 weeks (**Fig. 5e**). Similar to our data from *Cish*^*-/-*^ mice, there was no obvious accumulation of NK cells in mice that received g26-RNP CD34^+^ cells (**Fig.5f**), and NGS sequencing of the resulting NK cells showed a similar indel frequency to the starting g26-RNP CD34^+^ cord blood progenitors (**data not shown**) suggesting CIS is not playing a major role in human NK cell development and homeostasis. The NK cell population reconstituted from g26-RNP edited CD34^+^ cord blood progenitors phenocopied that of human peripheral blood NK cells as we have previously shown^31^ and did not differ from Mock-RNP controls (**data not shown**). We next interrogated the IL-15 receptor signaling of g26-RNP CD34^+^ cord blood progenitor-derived NK cells *in vitro*. We isolated human NK cells from the spleens of two g26-RNP and two Mock-RNP humanized mice and subjected them to an IL-15 proliferation assay *in vitro*. Similar to the data observed in g26-RNP primary human NK cells and g57-RNP SNK10 cells, the NK cells derived from g26-RNP humanized mice were able to proliferate significantly greater in response to IL-15 compare to their Mock-RNP counterparts (**Fig. 5g**). Furthermore, the indel frequency increased from 43% upon NK cell isolation from the g26-RNP humanized mouse spleen to 75% following the 7-day proliferation assay (**data not shown**). Taken together, these data demonstrate the efficiency of gene targeting in human NK cells and the validates the role of CIS as a potent negative regulator of IL-15 signaling in human NK cells.

**Figure 3.**
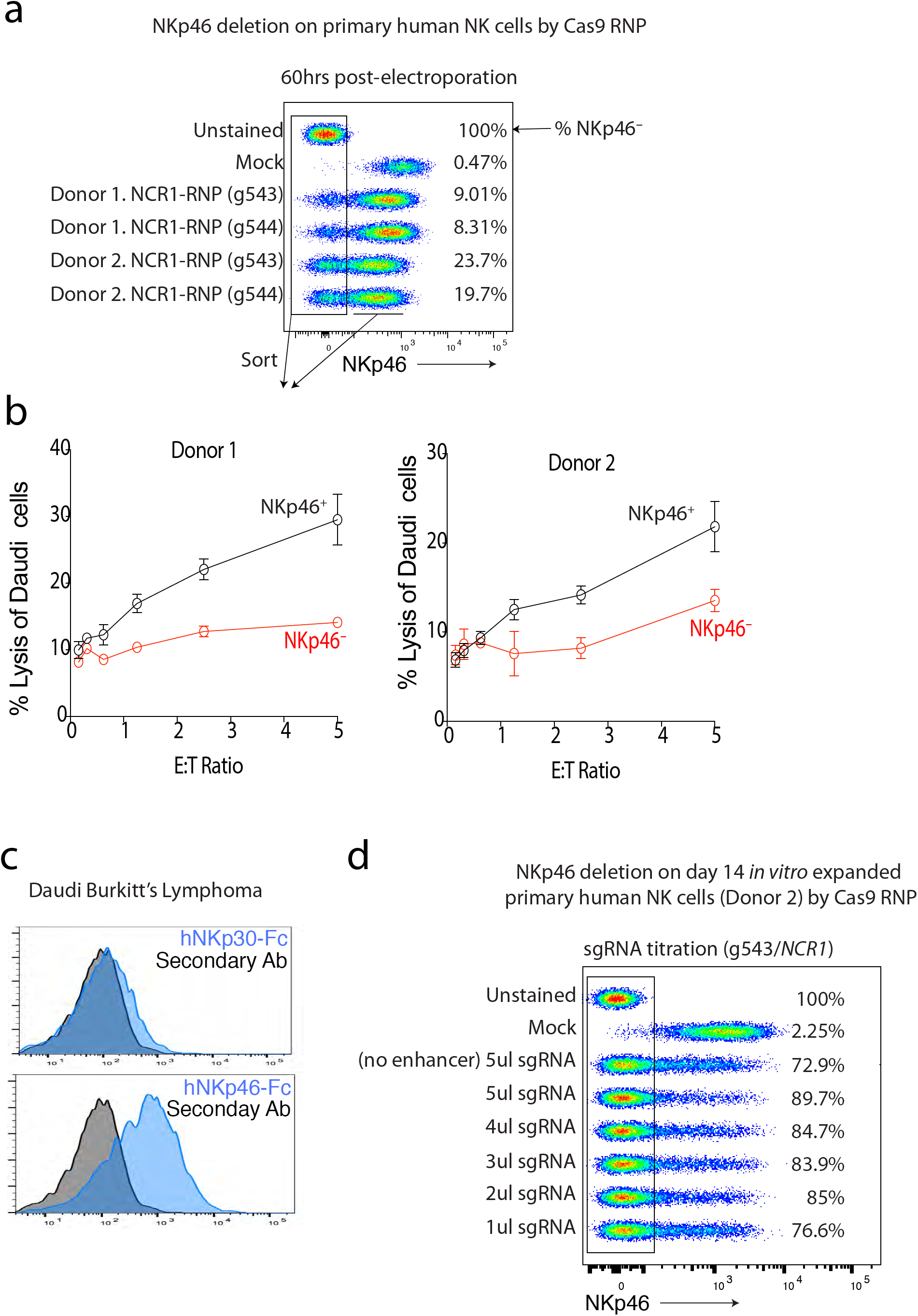
Deletion of *NCR1* using Cas9 RNP. (**a**) Deletion efficiency of *NCR1* 5-days post electroporation using two different guides shows some variability between donors. Data are representative of 4 healthy adult donors. (**b**) FACS-sorted NCR1^+^ and NCR1*-* primary human NK cells were used in a cytotoxicity assay against Daudi target cells. All data are shown. (**c**) Daudi target cells express high levels of putative NKp46 ligands as revealed by staining with an hNKp46-Fc. hNKp30-Fc served as a negative control. Data are representative of 2 experiments. (**d**) Deletion efficiency of *NCR1* on highly activated primary human NK cells (14 days post-expansion; donor 2 from **3a**) using a titration g543/*NCR1*. NKp46 staining was performed 60 hours post-electroporation. Percentages indicated *NCR1*-deleted (NKp46*-*) fraction. All data are shown from donor 2. Mock-RNP (electroporation of g543 alone) served as a negative control in all experiments.

**Figure 4.**
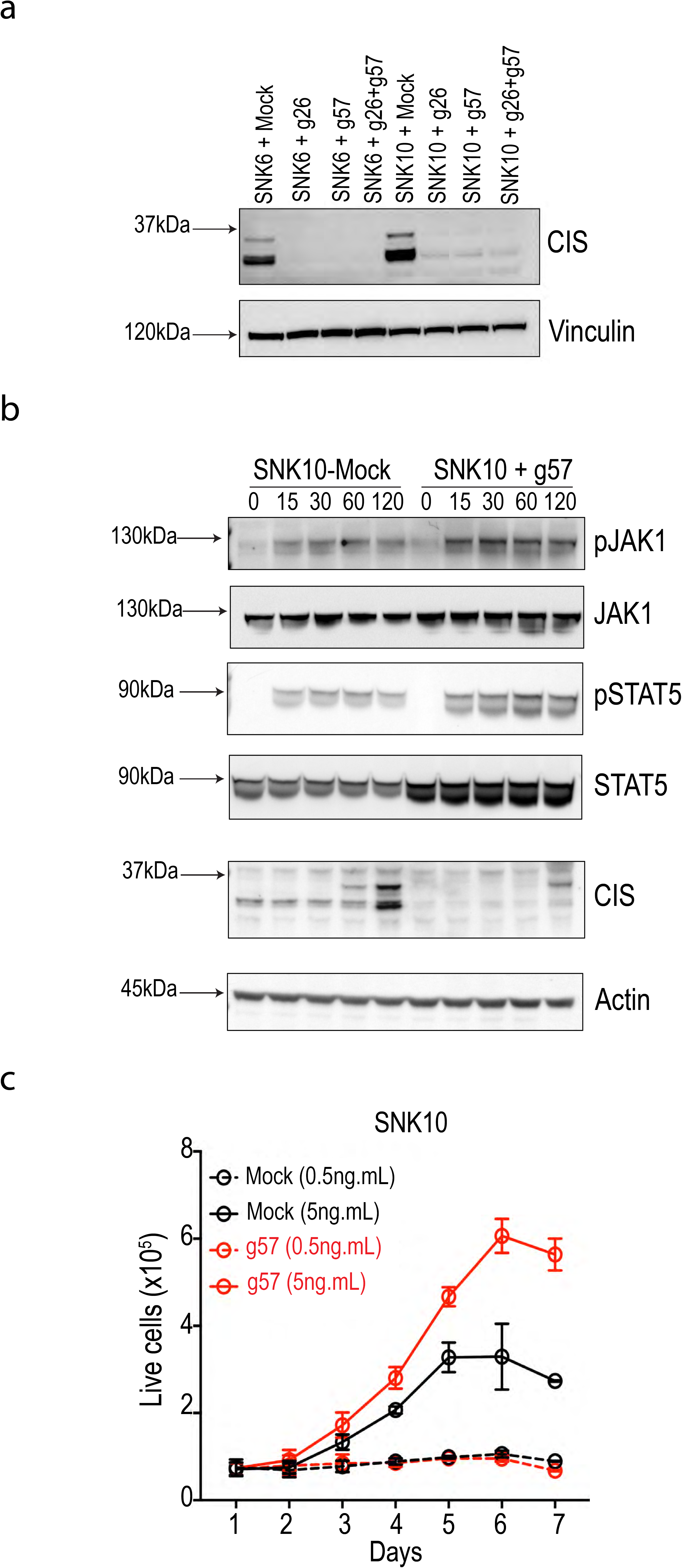
Deletion of *CISH* enhances cellular responses to IL-15. (**a**) Cas9 RNP-mediated deletion of CISH from SNK6/SNK10 NK cell lines using various guides alone or in combination. (**b**) CIS-deficient SNK10 cells show enhanced activation of the IL-15 signaling pathway relative to parental controls. (**c**) Proliferative response to IL-15 is enhanced in CIS-deficient SNK10 cells. Data are representative of 2 experiments.

**Figure 5.**
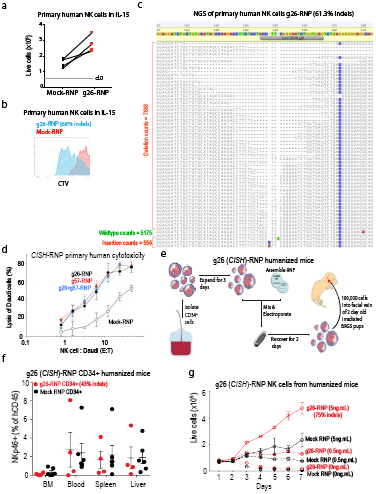
Deletion of CISH in primary human NK cells. NK cells were isolated from 4 healthy donors and electroporated with Cas9 RNP+g26(*CISH*) or Mock RNP (g26 alone). Proliferation was determined 5 days after electroporation by (**a**) seeding 50,000 cells/well in a 96-well plate and enumerating the number of live cells or (**b**) labeling cells with a division-tracking dye (CTV). *CISH* deletion efficiency was quantified for all donors and ranged from 45-80%, representative NGS results shown in (**c**). (**d**) *CISH*-edited primary human NK cells (g26, g57 or g26+g57) and Mock-RNP control NK cells from the same donor were tested for their cytotoxicity against Daudi B lymphoma cells *in vitro*. Data are representative of 2 donors (**e**) Workflow used to generate gene-edited human immune system (HIS) mice. (**f**) Frequency of human NK cells in the bone marrow (BM), spleen, liver and blood of HIS mice 12 weeks after reconstitution with WT (Mock RNP) or *CISH*-g26 edited CD34^+^ hematopoietic stem cells. (**g**) Proliferation of NK cells derived from the spleen of HIS mice (**e, f**) in various concentrations of IL-15, NGS revealed that CISH-indels were enriched from 43% to 75% after 7 days in culture.

## Discussion

Here we describe, for the first time, an effective approach for genetic manipulation of primary human NK cells using CRISPR-Cas9 RNP and demonstrate that our recent discovery of an NK cell checkpoint in mice is also a conserved and valid target in human NK cells. To date, there have been few examples of genetically-engineered primary human NK cells largely due to poor infection rates with viral gene-transfer systems and difficulties in culturing primary human cells to significant numbers. Recent advances in viral pseudotyping and antagonizing intracellular anti-viral pathways have improved infection rates and opened the door for application such as CAR-NK cell products^16^. A concern from lentiviral modification of human NK cells is mutagenesis caused by random insertion of viral genes into the genome given the high viral titers (>150 MOI) required for infection^17^. In addition, viral delivery and expression of gene-editing machinery require the target cells to be cultured for several days before the desired gene is edited, potentially losing the naïve phenotype of the target cell. A recent study showed that simply delivering plasmids that encode the gene editing machinery could efficiently target genes in T cells^24^, however our data suggest that even highly-activated NK cells are inefficient at transcribing the required genes from large plasmid templates to result in Cas9-mediated editing. It is worth noting that pMAX-GFP vector uses a CMV promoter and at 3486bp is considerably smaller than the Px458 (9300bp) which uses a Cbh promoter.

We therefore set out to establish whether direct delivery of Cas9-RNP complexes directly into freshly isolated human NK cells could rapidly and efficiently edit certain genes. An additional benefit of using NLS-tagged Cas9 RNP is that the delivery method does not depend upon successful nuclear delivery, as the complex can be easily transported into the nucleus^19^. Upon entry into the nucleus, the Cas9 RNP rapidly binds DNA and begins scanning for the regions complementary to the guide RNA. The kinetics of this editing process have been described to take between 2-12 hours^32^, meaning that primary human NK cells could be easily edited within a day to study their function.

The cancer immunotherapy revolution over the past 10 years has seen rising interest in targeting NK cells given their spontaneous ability to detect and kill stressed/transformed cells^6^. Furthermore, NK cells are emerging as key drivers of dendritic cell infiltration and optimal priming of IFN^+^ and TNF*a*^+^ CD8 T cells^3,4,10^. Yet, despite our best efforts, cancer immunotherapy drugs that exploit the innate anti-tumor functions of NK cells are yet to have a positive impact on cancer outcomes. This is in part due to the vast differences between mouse and human NK cell frequencies and phenotypes *in vivo* and due to the difficulty in validating if therapeutic targets derived from murine genetic knockout studies hold true for human NK cells. To this end, our CRISPR-Cas9 RNP approach detailed here will be transformative for NK cell immunotherapy drug development and basic understanding of human NK cell biology. CRISPR-Cas9 RNP now makes ease of rapid gene editing in primary human NK cells overcoming this current barrier to target pathway validation. To this end, we investigated two pathways known to regulate murine NK cell anti-tumor function in human NK cell lines and primary human NK cells. NKp46 is a transmembrane receptor that signaling via CD3ζ and FcεRIγ to recruit multiple ITAM-containing proteins. Cross-linking NKp46 with antibodies results in IFNγ production, which is augmented by IL-15 signaling^7^. NKp46-null mice display impair anti-metastatic responses *in vivo* and impaired cytotoxicity *in vitro* to tumor cells that bind NKp46-Fc and presumable express putative NKp46 ligands^27,33,34^. Using Cas9-RNP+g*Ncr1*, we were able to delete NKp46 on a subset of primary human NK cells and validate that NKp46 expression confers optimal NK cell cytotoxicity towards NKp46-ligand expressing Daudi B lymphoma cells. Deletion of genes for cell-surface proteins was thus clearly feasible, and the ability to FACS sort for gene-edited cells makes cell surface proteins practical targets for validation using this method. We next addressed the feasibility to delete the intracellular immunotherapy drug target CIS using Cas9-RNP+g*Cish*. *Cish* has been published by multiple groups to limit IL-15 and TCR signaling in murine NK and T cells, ultimately dampening anti-tumor responses^7,35,36^. *Cish* was efficiently deleted in human NK cells (45-80% indels in primary cells; 85-93% indels in NK cell lines) yet the inability to sort on *Cish*-null cells meant functional assays were run on pools of deleted cells. Despite the incomplete absence of CIS protein, Cas9-RNP+gCish edited cells behaved as predicted from mouse studies in that they were hyper-sensitive to IL-15 *in vitro* and had augmented cytotoxicity towards an allogeneic tumor line *in vitro*. Much resources have been placed on optimizing NK cells as an adoptive off-the-shelf cellular immunotherapy. These products derive from numerous precursor cell populations, including induced pluripotent stem cells, placental NK cells and cord-blood progenitors^37^. Given that deletion of *Cish* in mature human NK cells phenocopies the IL-15 hyper-responsiveness data from mouse^7^, it is expected that CRISPR deletion of *Cish* in human adoptive NK cell therapy products would augment their anti-tumor function as previously shown for murine *Cish*^-/-^ NK cells.

Thus, while primary human T cells can be genetically modified by several approaches including lenti/retrovirus transduction, transient transfection and Cas9-RNP^13, 24^, the latter approach is the only viable approach for large-scale genetic modification of primary human NK cells. With further optimization of guide sequences, recombinant Cas9 source and multiplexing of distinct guides, it is likely that most genes will be able to be deleted at a high enough efficiency to concinving conclude their role in human NK cells. Our detailed technical report of Cas9-RNP in human NK cells lends itself to immunotherapy drug discovery including rapid gene-editing of patient-derived or allo-NK cellular therapy products meaning undruggable pathways can be efficiently targeted for personalized immunotherapy.

## Acknowledgements

This work was supported by project grants from the National Health and Medical Research Council (NHMRC) of Australia (1124907, 1124784, 1049407, 1066770, 1057852, 1027472) and a fellowship (1124788) to N.D.H as well as an NHMRC Independent Research Institute Infrastructure Support scheme grant and a Victorian State Government Operational Infrastructure Scheme grant. N.D.H. is a recipient of Research Grants from the Harry J Lloyd Charitable Trust (USA), Melanoma Research Alliance (USA), Tour de Cure (AUS), the Ian Potter Foundation (AUS), Cancer Council of Victoria (1145730) and a CLIP grant from Cancer Research Institute (USA).

## Authorship Contributions

JR, ES and NDH designed and performed experiments and interpreted all data. JR and NDH wrote the paper.

## Disclosure of Conflicts of Interest

JR and NDH are co-founders and share-holders of oNKo-Innate Pty Ltd. NDH has a collaborative research agreement with Servier.

